# Microbiome Modulation Uncouples Efficacy and Toxicity Induced by Programmed Death-1/Programmed Death-Ligand1 Blockade

**DOI:** 10.1101/2024.05.08.590282

**Authors:** Laura Lucia Cogrossi, Paola Zordan, Matteo Grioni, Anna Tosi, Nathalie Rizzo, Anna Policastro, Benedetta Mattorre, Marco Lorenzoni, Greta Meregalli, Sofia Sisti, Francesca Sanvito, Marta Chesi, Leif Bergsagel, Nicola Clementi, Antonio Rosato, Matteo Bellone

## Abstract

While asymptomatic smoldering multiple myeloma (SMM) holds an overall risk of progression to multiple myeloma (MM) at 10% per year, only active surveillance is offered to most patients affected by SMM, which leaves them in anxiety and frustration. Intestinal microbiota and gut-born T helper 17 (Th17) lymphocytes may act as drivers of MM evolution. In transgenic Vk*MYC mice developing *de novo* MM, which invariably evolves from Early-MM that mimics SMM to full-blown Late-MM, we investigated the impact of gut microbiota modulation on disease progression and susceptibility to immune checkpoint blockade (ICB). We report that administering the human commensal *Prevotella melaninogenica* to mice affected by Early-MM significantly delayed evolution to Late-MM. Mechanistically, treatment with *P. melaninogenica* induced increased production of short chain fatty acids. Butyrate prevented skew of dendritic cells towards a pro-Th17 phenotype and treated mice accumulated less disease induced Th17 cells in their bone marrow. *P. melaninogenica* also synergized with anti-PD-L1 antibodies by restraining Th17 cell expansion while unleashing ICB-induced full effector CD8^+^ T cells, eventually blocking progression to full-blown disease. Similar results were obtained in mice challenged with bortezomib-resistant Vk*MYC tumor cells, a model of more aggressive MM. When mice were exposed to imiquimod to mimic ICB-associated psoriasis-like lesions, *P. melaninogenica* ameliorated skin lesions caused by ICB. Thus, modulation of the gut microbiota with *P. melaninogenica* might represent a treatment for patients affected by SMM and would allow fully exploiting the antitumor potential of ICB in plasma cell dyscrasias.

**Key points:**

Administration of the human commensal *Prevotella melaninogenica* to Vk*MYC mice delayed evolution to symptomatic multiple myeloma;
*P. melaninogenica* therapeutically synergized with PD-1/PD-L1 blockade also limiting immune-related adverse events.

## Introduction

Multiple myeloma (MM) is a treatable but incurable B cell neoplasm characterized by the accumulation of clonal plasma cells within the bone marrow (BM), anemia, hypercalcemia, renal insufficiency, and bone lesions^1^. Smoldering multiple myeloma (SMM) is an asymptomatic, and in principle curable phase that may anticipate MM^2^. Likely because SMM is a heterogeneous disease and its pathobiology is poorly understood, treatment is restricted to few high-risk patients, whereas active surveillance is recommended to all the others^3^. Considering that SMM has an overall risk of progression to MM at 10% per year^4^, active surveillance leaves many SMM patients in anxiety and frustration.

The intestinal microbiota may represent one tumor cell-extrinsic mechanism driving SMM-to-MM transition^5,6^. In transgenic Vk*MYC mice developing *de novo* MM^7^, *Prevotella heparinolytica*, Gram-negative bacteria belonging to the human intestinal microbiota^8^, promoted the differentiation of gut-born Th17 cells that migrated to the BM where they favored neoplastic plasma cell expansion, and evolution from asymptomatic disease mimicking SMM (i.e. Early MM; ref.^9^) to symptomatic MM (Late-MM; ref.^10^). Conversely *P. melaninogenica*, which correlate with better prognosis in humans affected by autoimmune diseases^11^, restrained Th17 cell expansion and prolonged animal survival. Mechanistically, IL-17 induced STAT3 phosphorylation in neoplastic plasma cells and activated eosinophils. Blocking the IL-17 pathway delayed disease progression in Vk*MYC mice^10^. Similarly, in MM patients affected by SMM, higher levels of BM IL-17 predicted faster disease progression^10^. Altogether, these premises suggest that microbiota modulation would represent a safe SMM treatment.

The intestinal microbiome has also gained momentum in the context of cancer immunotherapy, and, because the gut microbiota can empower immune checkpoint blockade (ICB)^12–14^, clinical trials of oral supplementation with probiotics in combination to ICB are ongoing (NCT03358511, NCT03829111, NCT05032014, NCT05094167, NCT05865730).

T cells in MM patients express PD-1 and are functionally exhausted/senescent^15^, and PD-1-expressing T cells are enriched in the BM of SMM patients who progressed to active MM^16^. ICB targeting PD-1/PD-L1 have been tested in MM also supported by the evidence that PD-L1 is selectively expressed on neoplastic plasma cells^17^. PD-L1 may also act on plasma cells as oncogenic protein, and its expression correlates with disease progression and relapse^18^. However, all trials combining anti-PD-1/PD-L1 antibodies, dexamethasone and immune modulatory drugs in MM were halted by US FDA because of unfavorable benefit-risk profile and life-threatening immune-related adverse events (IrAEs)^19^. Because defined microbiota signatures have been associated with IrAEs ^20–24^, which can be induced by IL-17-producing cells ^25,26^, probiotic administration might represent a valid strategy to limit IrAEs while improving anti-tumor immunity.

Here, we show that modulation of the gut microbiota through administration of *P. melaninogenica* is a disease-modifying treatment *per se*. Additionally, when given in combination with anti-PD-L1 antibodies, it substantially modifies the trajectory of SMM-to-MM progression in Vk*MYC mice and limit ICB-related skin toxicity, thus fully exploiting the antitumor potential of ICB in plasma cell dyscrasias.

## Methods

### Mice

All mice were on a C57BL/6 background. Wild type (WT) mice were purchased from Charles River Laboratories, Calco IT. In Vk*MYC transgenic mice^7^ the activation of MYC, whose locus is rearranged in half human MM tumors including SMM^27^, occurs sporadically through the somatic hypermutation process in germinal center B cells. All mice were bred and maintained in pathogen-free or in conventional animal facilities. Animal procedures were approved by the institutional animal ethics committee (Italian Ministry of Health authorization No. 208/2023-PR).

### Bacteria cultivation

*P. melaninogenica* DSM 7089 and *P. heparinolytica* DSM 23917 (DSMZ, Germany) were cultured in chocolate agar plates at 37 °C in AnaeroJar 2.5 L Jar System (OXOID) and using AnaeroGen 2.5L (Thermo Scientific) to generate an anaerobic atmosphere.

### Bacteria and antibody treatment *in vivo*

Early-MM mice^9^ were pretreated for one week with antibiotic mix containing Ampicillin (1g/L, Sigma-Aldrich), Vancomycin (0.5g/L, Sigma-Aldrich), Neomycin (1g/L, Sigma-Aldrich) dissolved in drinking water and Metronidazole (0.2mg/mouse, Sigma-Aldrich) via gavage three times. 48h after antibiotic withdrawal, *P. melaninogenica* was administered via gavage for three consecutive days/week until the end of the experiment. Control mice received gavage with saline solution (PBS). Anti-PD-L1 monoclonal (B7-H1, clone 10F.9G2, BioxCell; 200 µg/dose) or isotype antibodies (Rat IgG2b) were injected i.p. on day 0, 5, 8, and every three weeks until the end of the experiment. Every three weeks mice were bled for monoclonal (M) protein (M-spike) quantification as described in Supplementary Methods section. Mice were sacrificed at Late-MM or at the end of observation time. Alternatively, two weeks before tumor cell injection (I.V. 1×10^6^ Vk12598 cells obtained from one Late-MM mouse; t-Vk*MYC^7^), antibiotic pretreated WT mice were subjected to gavage with *P. melaninogenica*, or *P. heparinolytica* or PBS and received anti-PD-L1/IgG three times/week. t-Vk*MYC mice were weekly bled for M-spike quantification. For immunological experiments, mice were sacrificed at M-spike appearance, and harvested organs were analyzed as described in Supplementary Methods section.

### Butyrate treatment

Sodium butyrate (100mM; Sigma-Aldrich) was dissolved in drinking water of WT mice two weeks before tumor injection (t-Vk*MYC mice). Solution was refreshed b.i.w. Control mice received plain water.

### Imiquimod-induced skin inflammation

Antibiotic-pretreated mice received oral gavage of *P. melaninogenica* or PBS until the end of the experiment. Three doses of anti-PD-L1 or IgG monoclonal antibodies (of 200µg each) were injected i.p. every 4 days. Subsequentially, mice received a topical dose of 62,5mg of Imiquimod cream (5%) (Aldara; 3M Pharmaceuticals) on the shaved back skin for 7 consecutive days. Each dose corresponded to 3,125mg of active compound^28^. On day 7, organs were collected for histopathological evaluation and flow cytometry analyses.

### Multiplex immunohistochemistry (mIHC)

The Tyramide Signal Amplification (TSA)-based Opal method (Akoya Biosciences, Marlborough, MA, USA) was used for mIHC staining on the Leica BOND RX automated immunostainer (Leica Biosystems, Wetzlar, Germania), as previously described^29^. All stains were performed under optimized conditions. The spectral DAPI (Akoya) was used as nuclear counterstain, and slides were mounted in ProLong Diamond Anti-fade Mountant (Life Technologies, Waltham, MA, USA). An unstained tissue section was used to subtract the tissue autofluorescence from multiplex-stained slides. Imaging and data analysis are described in Supplementary Method section. Primary antibodies are listed in Table S2.

### Histopathological evaluation and scoring severity of skin inflammation

Histopathological evaluation was performed in blind by two expert pathologists on 2 H&E-stained slides of skin for mouse. Hyperkeratosis, acanthosis, spongiosis, and inflammatory infiltrate were scored independently on a scale from 0 to 4: 0, none; 1, slight; 2, moderate; 3, marked; 4, very marked.

### *In vitro* induction of mouse BM-derived dendritic cells

BM cells collected from long bone flushing were cultured for 6 days in IMDM (Lonza Bioscience) with 10% FCS (Fetal Calf Serum), GM-CSF (25ng/mL; PeproTech, Inc.) and IL-4 (5ng/mL; PeproTech). DCs were induced to overnight maturation with 1mg/mL heat-inactivated *P.heparinolytica*, *P. melaninogenica* or 1µg/mL LPS from *E.coli*.

### *In vitro* induction of human monocyte-derived DCs

CD14^+^ monocytes were isolated by magnetic sorting (Miltenyi Biotec) from healthy donor’s peripheral blood mononuclear cells (PBMCs) and cultured in RPMI (Lonza Bioscience) with 10% FCS, GM-CSF (50ng/ml; Peprotech, Inc) and IL-4 (25ng/ml; Peprotech, Inc). DC maturation was induced as described above.

### Mouse Th17 induction *in vitro*

Splenocytes from OTII mice^30^ were pulsed with 50µg/10^6^ cells of OVA_323–339_ peptide for 1h and plated at 10:1 ratio with mature BMDCs. Coculture was maintained for 5 days with TGF-β (2ng/mL; Peprotech, Inc.), IL-6 (10ng/mL; Peprotech, Inc.), anti-mouse IFNγ (10µg/mL; Biolegend, Inc) and anti-mouse IL-4 (10µg/mL; Biolegend, Inc).

### Human Th17 induction *in vitro*

The PBMC CD14^-^ fraction was plated at 10:1 ratio with HLA-matched MoDCs preconditioned with bacteria as described above. Dynabeads™ Human T-Activator CD3/CD28 (ThermoFisher Scientific) were added to the culture. Coculture was maintained for 6 days in the presence of TGF-β (1,12ng/mL Peprotech, Inc.), IL-23 (15ng/mL; Peprotech, Inc), anti-human IFNγ (1 μg/mL; Biolegend, Inc) and anti-human IL-4 (2μg/ml; Biolegend, Inc).

### Fecal short-chain fatty acid quantification

Feces were resuspended in buffer (PBS pH7.4,10%D2O and 0.01% NaN3; ratio 1:10 mg feces:μL buffer). Samples were centrifuged for 5’ at 10.000rpm and supernatants were collected. 50μM of 3-(Trimethylsilyl)propane-1-sulfonic acid, as internal chemical shift reference, was added to samples and NMR spectra acquired. NMR spectra were recorded at 298K on a Bruker Avance 600MHz spectrometer equipped with triple resonance cryoprobe. 1D-1H-NMR spectra (noesypr1d) were recorded with an acquisition time of 2s, 128 transients and a relaxation delay of 6s. Spectral window was set to 12ppm. Spectra were processed with zero filling to 132k points, and apodized with an unshifted Gaussian and a 1 Hz line broadening exponential using Mnova 14.0 (Mestrelab Research).

### Statistical analysis

Sample size was chosen considering the means of target values between experimental and control groups, standard error and statistical analyses used. Animals were matched for age and sex. Randomization was performed for *in vivo* experiments. Data were analyzed with GraphPad Prism version 9. Data are presented as mean ± standard deviation of the mean, individual values as scatter plot with column bar graphs. Differences were considered significant when P<0.05. N values represent biological replicates. Survival curves were compared using the log-rank test (Mantel–Cox). All the statistics and reproducibility are reported in the figure legend.

## Results

### *P. melaninogenica* either alone or in combination with anti-PD-L1 antibodies delays progression from Early-MM to Late-MM

To investigate the effect of *P. melaninogenica* on SMM-to-MM progression, we set up a preclinical trial enrolling Vk*MYC mice when they reached the phase of asymptomatic Early-MM [i.e., measurable M-spike < 6% which corresponds to gamma/albumin ratios ranging between 0.15 and 0.4 (Fig. S1A; ref.^9^). This model is relevant because Early-MM mice, if left untreated, invariably evolve to symptomatic Late-MM, thus mimicking those 10% SMM patients that each year progress to MM^10^. At enrollment, all mice displayed similar levels of M-spike (i.e., gamma globulins/albumin ratio; Fig. S1B) and normal hemoglobin (Hb) levels (Fig. S1C). To favor acceptance of the commensal, mice received broad spectrum antibiotics in the week before being randomly assigned to either receive *P. melaninogenica* or vehicle (PBS; Fig. 1A). Mice reached Late-MM when their M-spike was ≥ 6% (i.e., gamma/albumin ratio ranging between 0.35 and 0.75 Fig. S1A; ref.^9^). Hb drop was also taken as indication of Late-MM ^9^ (Fig. 1B and C). At week 14, M-spike levels in *P. melaninogenica*-treated mice were halved compared to PBS-treated mice (Fig. 1D). By that time, 50% of mice in the *P. melaninogenica*-treated group evolved to Late-MM compared to 90% in the PBS group, and overall progression to advanced MM was delayed by *P. melaninogenica* (Fig. 1E). Thus, a probiotic treatment *per se* was sufficient to delay disease evolution in Early-MM mice.

**Figure 1.**
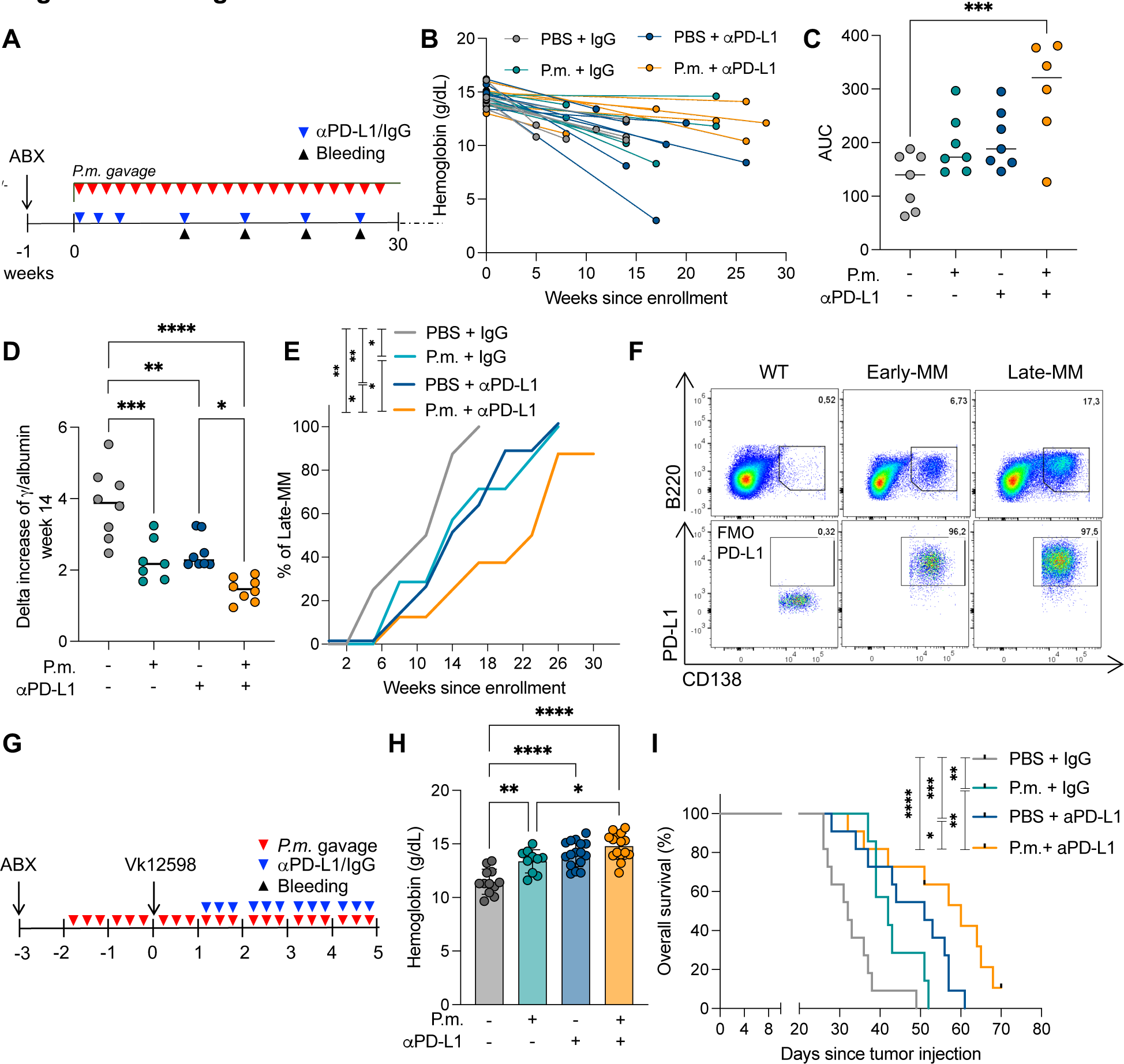
Combination of probiotic *P. melaninogenica* and anti-PD-L1 in Early-MM mice delays progression to Late-MM and prolongs survival in bortezomib-resistant MM. **A.** Treatment schedule for experiments B-E: Early-MM mice were enrolled, pretreated with antibiotics (ABX) and randomly assigned to the following treatments: Vehicle (PBS) + Isotype control (IgG), (n=8); anti-(α)PD-L1 + IgG (n=8); PBS + P.m. (n=7); P.m.+ αPD-L1 (n=8). M-spike levels and anemia were measured in the peripheral blood (Bleeding). **B**. Hemoglobin quantification in the peripheral blood of each mouse described in A at enrollment and at the time of Late-MM progression. **C**. Quantification of the area under the curve (AUC) for each mouse represented in B. ***P<0.001 by One-Way ANOVA. **D.** Delta increase of γ/albumin ratio for each mouse 14 weeks after enrollment. Error bars represents ± SD. *P<0.05, **P<0.01, ***P<0.001, ****P<0.0001 by One-Way ANOVA. **E.** Percentage of Early-MM Vk*MYC mice progressing to Late-MM overtime. *P<0.05, **P<0.01 by RM-One-way ANOVA. **F.** Representative flow cytometry plots of PD-L1 expression gated on BM plasma cells (identified as B220^-^CD138^+^ cells in CD45^+^ cells) from WT, Early-MM, Late-MM mice. **G.** Treatment schedule for C57BL/6 mice injected with Vk12598 MM cells (t-Vk*MYC) conditioned with P.m. or vehicle (PBS) and receiving αPD-L1 or IgG. M-spike levels and anemia were measured weekly in the peripheral blood. **H.** Hemoglobin quantification from mice treated as in G, 4 weeks after tumor injection. Each dot represents one mouse. Error bars represents ± SD. *P<0.05, ***P<0.001, ****P<0.0001. One-Way ANOVA. **I.** Overall survival of t-Vk*MYC mice described in G. PBS+IgG n=15, P.m.+IgG n=8, PBS+antiPD-L1 n=15, P.m.+antiPD-L1 n=15. Log-rank (Mantel–Cox) test. *P<0.05, **P<0.01, ***P<0.001, ****P<0.0001. Data are pooled from three independent experiments.

Because the gut microbiota also contributes to ICB efficacy^12–14^ and neoplastic plasma cells both in humans^17^ and in Early-MM mice express PD-L1 (Fig. 1F), we treated Early-MM mice with anti-PD-L1 antibodies alone or in combination with *P. melaninogenica*. Early-MM mice treated with anti-PD-L1 experienced delayed progression to Late-MM (Fig. 1D,E). Addition of *P. melaninogenica* to anti-PD-L1 further delayed disease evolution (Fig. 1D,E) and, when all Vk*MYC mice in the control group had evolved to Late-MM, more than 60% of the mice receiving the combined treatment were still in the Early-MM phase (Fig. 1E). At the end of the observation period, 20% of mice in the combination group had not evolved to Late-MM (Fig. 1E). We did not notice any substantial difference in response to therapy between male and female mice (Fig. S1D-G). Additionally, most of the mice receiving the combined treatment experienced limited Hb drop (Fig. 1B), as confirmed by the comparison of the area under the curve (AUC) between this and the PBS group (Fig. 1C).

We also assessed the therapeutic efficacy of *P. melaninogenica* and ICB in mice challenged with Vk12598 cells (t-Vk*MYC; Fig. 1G), a model that mimics bortezomib-resistant MM and already predicted therapeutic efficacy in humans^31^. Four weeks after tumor injection, all control mice were anemic (i.e., Hb ≤13 g/dL; Fig. 1H; ref.^7^) and 80% of them reached humanitarian end point 35 days after tumor injection (Fig. 1I). *P. melaninogenica* or anti-PD-L1 given as single treatment limited anemia (Fig. 1H) and prolonged mouse survival (Fig. 1I). Importantly, the combination of *P. melaninogenica* and anti-PDL-1 provided additional survival advantage (Fig. 1I) and further contained anemia (Fig. 1G). Thus, *P. melaninogenica* and ICB synergize in limiting aggressiveness of MM.

### *P. melaninogenica* limits Th17 cell expansion while unleashing ICB-induced full effector T cells

Because IL-17 has been associated with faster SMM-to-MM progression and in Vk*MYC mice IL-17-producing cells are modulated by the gut microbiota^10^, we quantified IL-17-producing cells in the intestine and in the BM of treated Vk*MYC mice at the end of the observation period. Except for one mouse in the combined treatment group that never progressed to Late-MM, most of the mice were sacrificed at similar levels of M-spike (Fig. S1H), which is indicative of BM infiltration by neoplastic plasma cells^9^. Despite comparable tumor burden and total CD4^+^ T cells in both Peyer’s patches (PPs; Fig. S2A,B) and BM (Fig. S2F,G) of mice receiving either treatment, Th17 cells were less represented in the PPs (Fig. 2A,B and Fig. S2D) and in the BM of *P. melaninogenica*-treated mice (Fig. 2C,D and Fig. S2I). Strikingly, this was also true for mice exposed to the combined treatment (Fig 2A-D and Fig. S2D,I). Similar results were obtained in t-Vk*MYC mice exposed to single or combined treatments (Fig. 2 SK-N). Hence, *P. melaninogenica* contained disease-induced expansion of Th17 cells even under ICB treatment.

**Figure 2.**
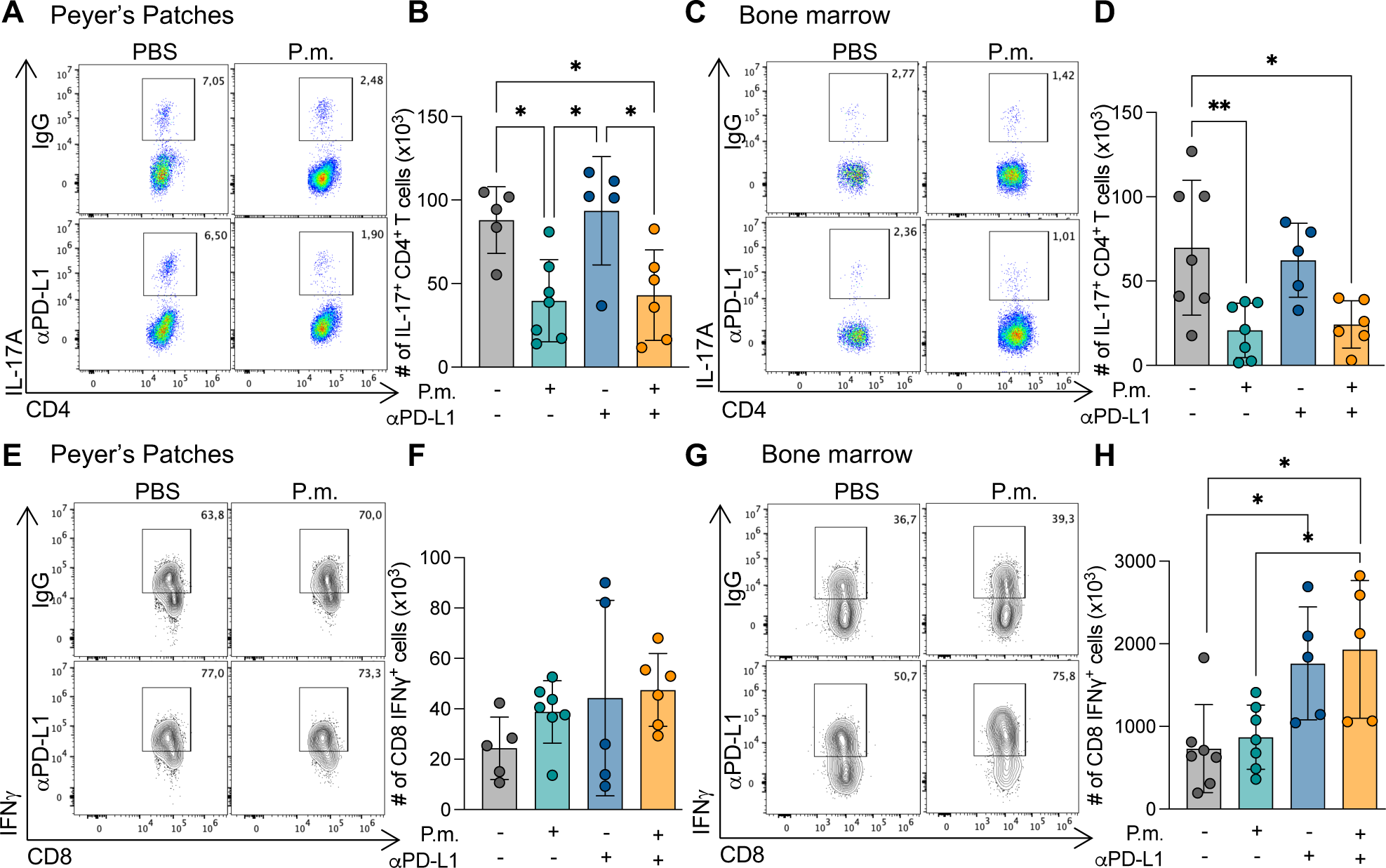
*P.melaninogenica* limits Th17 cell expansion fully unleashing the cytotoxic response elicited by anti-PD-L1. **A.** representative flow cytometry plots for IL-17 production gated in CD3^+^CD4^+^ T cells (Th17 cells) in the PPs of animals described in fig. 1A-E. **B**. Quantification of Th17 cells in the PPs from flow cytometry plots represented in A. **C.** Representative flow cytometry plots for Th17 cells in the BM of the same animals. **D.** Quantification of Th17 cells in the BM from flow cytometry plots represented in **C**. **E.** Representative flow cytometry plots for IFNγ production gated in CD3^+^CD8^+^ T cells in the PPs of animals described in fig. 1A-E. **F.** Quantification of IFNγ producing CD8 T cells in the PPs from flow cytometry plots represented in E. **G.** Representative flow cytometry plots for IFNγ production gated in CD3^+^CD8^+^ T cells in the BM of the same animals. **H.** Quantification of IFNγ−producing CD8^+^ T cells in the BM from flow cytometry plots represented in F. Error bars represents ± SD. *P<0.05, **P<0.01 One-Way ANOVA.

ICB is expected to rejuvenate CD8-mediated anti-tumor immunity^15^. Despite comparable total CD8^+^ T cells in both PPs (Fig. S2A,C) and BM (Fig. S2F,H) of mice receiving either treatment, much more IFNγ-producing CD8^+^ T cells infiltrated the BM of mice treated with ICB (Fig. 2G,H and Fig. S2E). ICB did not expand IFNγ-producing CD8 T cells in the PPs (Fig. 2 E,F and Fig. S2E). This potent CD8-mediated immune response was unaffected in mice also treated with *P. melaninogenica* (Fig. 2E-H), showing that the effect of *P. melaninogenica* substantially and persistently extends beyond the intestinal tract to the BM, where it reduces the expansion of pathogenic Th17 cells without dampening the antitumor response elicited by anti-PD-L1.

### *P. melaninogenica* restrains IL-17-mediated skin toxicity induced by ICB

Severe IrAEs in patients receiving ICB in combination immunomodulatory drugs^19^ precluded further investigation. The microbiota affects IrAEs especially in the gastrointestinal tract and the skin^32^, and IL-17 blocking antibodies can treat ICB-induced psoriasis^25^. Having provided evidence that *P. melaninogenica* contained Th17 cell expansion even under ICB, we asked if it also restrained IL-17 and IL-17-producing cells that ignite dermatological toxicity^33^. Because anti-PD-L1 does not induce skin toxicity in our models, we applied imiquimod to anti-PD-L1-treated mice to model ICB-induced psoriasis-like lesions^28^. Imiquimod-induced skin reaction is mediated by IL-17^34^ and the inhibition of PD-1 signaling in mice enhances IL-17-mediated psoriatic dermatitis^35^. Similarly, in anti-PD-L1-treated mice, Imiquimod generated psoriasis-like lesions characterized by parakeratosis, spongiosis and microabscesses that were moderated in mice also receiving *P. melaninogenica* (Fig. 3A). Histopathological evaluation of the skin sections confirmed that *P. melaninogenica* reduced inflammation (Fig. 3B), acanthosis (Fig. 3C) and hyperkeratosis (Fig.3D) in anti-PD-L1 treated mice. Additionally, mIHC (Fig. 3E,H and Fig. S3) showed a trend towards reduced density of IL-17^+^ cells (Fig. S3A,B) and reduced gut-born (i.e., α4β7^+^) cells in the skin of mice receiving both *P. melaninogenica* and anti-PD-L1 than imiquimod/anti-PD-L1 treated mice (Fig. 3F). The latter data suggest restrained migration of cells from the gut into the skin of mice receiving *P. melaninogenica*. CD11c^+^ (Fig. 3G) and CD11c^+^IFNγ^+^ cells (Fig. 3H) were also less represented and tended to be more distant from CD4^+^ and CD8^+^ T cells in *P. melaninogenica*-treated mice (Fig. S3C), suggesting that *P. melaninogenica* also impacted antigen presenting cell (APC) migration and function. Th17 cells accumulated in skin-draining lymph nodes of treated mice, which were restrained by *P. melaninogenica* (Fig. 3I). IFNγ-producing cells remained unaltered (Fig. 3J). Thus, *P. melaninogenica* uncoupled efficacy and toxicity of ICB.

**Figure 3.**
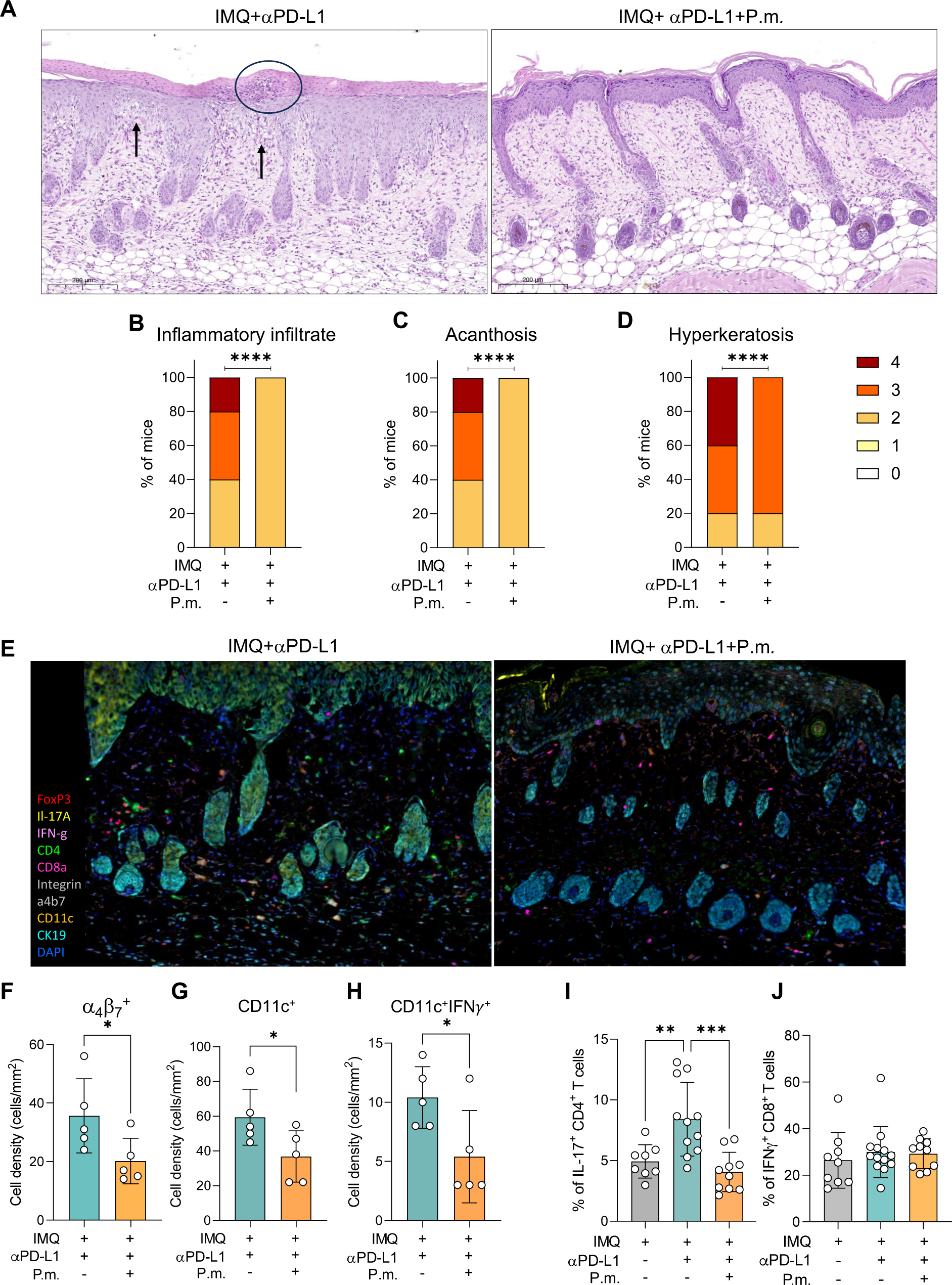
*P. melaninogenica* restrains IL-17-mediated dermatological toxicity upon ICB treatment. **A.** Representative images of skin histopathology (H&E staining) from mice treated with imiquimod (IMQ) + αPD-L1 in the absence (left) or presence (right) of P.m. Magnification 15X. Arrow indicates spongiosis, circle indicates microabscess. **B-D.** Stacked bar plot showing the percentage of mice according to the histopathological score for inflammatory infiltrate (**B**), Acanthosis (**C**), and hyperkeratosis (**D**) performed on 2 H&E stained slides of skin for each mouse. n=5mice/group. ****P<0.0001 Chi-square test. **E**. Representative multiplex IHC images from mice treated with IMQ + anti-PD-L1 in the absence (**left**) or presence (**right**) of P.m. administration. Magnification 20X. **F-H**. Cell density per slide of Integrin α4β7^+^ cells **(F**), CD11c^+^ cells (**G**) and IFNγ^+^CD11c^+^ cells (**H**). Each dot represents one mouse. Error bars represents ± SD. *P<0.05 by Unpaired Student’s t-test. **I-J.** flow cytometry staining on the skin-draining lymph nodes (inguinal) of the same mice. Data show the percentage of IL-17^+^ cells gated in CD4^+^ T cell (**I**) and IFNg^+^ cells gated in CD8^+^ T cells (**J**). Each dot represents one mouse. Error bars represents ± SD. *P<0.05 by One-Way ANOVA.

### *P. melaninogenica* modulates DC function to restrain Th17 expansion

Because gut microbiota conditioning by *P. melaninogenica* associated with reduced skin-infiltrating CD11c^+^ cells, we focused on DCs that are master regulators of Th skew^36^. Upon treatment of t-Vk*MYC mice with either one of the two *Prevotella* strains (Fig. 4A), more Th17 cells (Fig. 4B) and more DCs producing IL-6 and IL-1β (Fig. 4C) were found in the BM of mice receiving *P. heparinolytica*. IL-6 and IL-1β, together with IL-23, are involved in the differentiation of naïve T cells towards Th17 cells^37–39^, and the amount of IL-6 is increased in the BM of Vk*MYC mice, where it synergizes with IL-17 to promote neoplastic plasma cell survival^10^. Indeed, more DCs generated from BM precursors of WT mice produced IL-1β (Fig. 4D) when conditioned with *P. heparinolytica*, and, more importantly, skewed more naïve OTII cells ^30^ towards Th17 (Fig. 4E). Additionally, more cytokine-producing human monocyte derived DCs (hMoDCs) were found in *P. heparinolytica*-conditioned samples (Fig. 4F), and more naïve T cells became Th17 cells in the presence of *P. heparinolytica*-conditioned hMoDCs than *P. melaninogenica*-conditioned DCs (Fig 4G). Altogether, these findings support a mechanism by which *Prevotellae* modulate the immune response by conditioning DCs.

**Figure 4.**
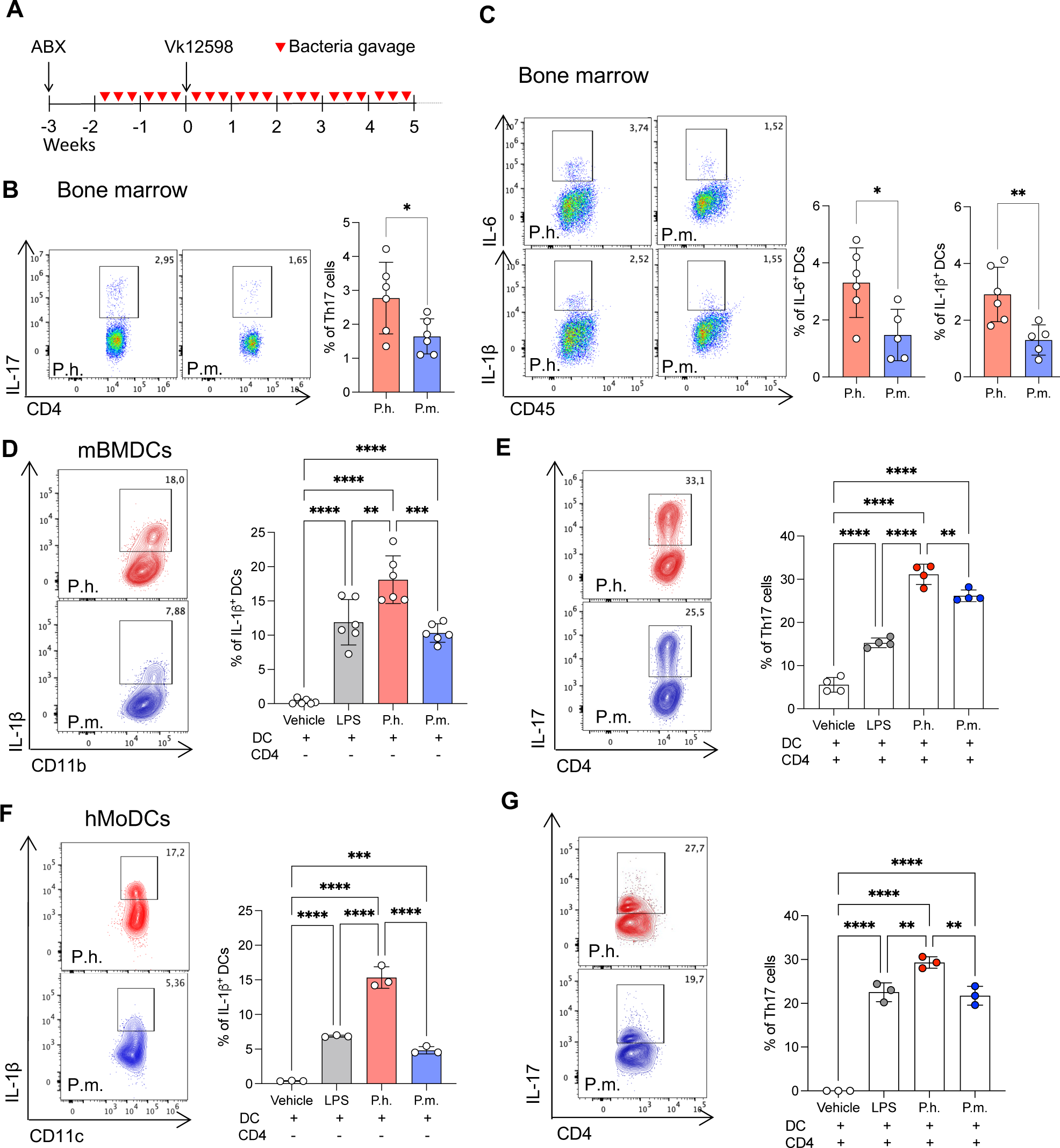
*P. melaninogenica* modulates mouse and human DC function to restrain the expansion of Th17 cells. **A.** Treatment schedule for t-Vk*MYC mice conditioned with P.m. or P.h. and sacrificed at the appearance of M-spike. **B-C**. immunological analysis performed by flow cytometry on mice described in A. **B**. percentage of Th17 cells gated in CD4^+^ T cells in the BM. **C.** Percentage of BM DCs (gated as CD11c^+^ cells) producing IL-6 and IL-1β. Each dot represents one mouse. Error bars represents ± SD. *P<0.05 Unpaired Student’s t-test. **D**. Percentage of mouse BMDCs producing IL-1β upon overnight stimulation with LPS from *E.coli* (1μg/mL), heat-inactivated P.h. or P.m. (1mg/mL). Vehicle: Medium. **E**. Percentage of Th17 cells (identified as IL-17^+^ cells in CD3^+^CD4^+^ cells) obtained upon coculture of muse OT-II cells with BMDCs from D. Coculture was maintained for 5 days under suboptimal Th17-polarizing condition (see methods section). **F.** Percentage of human MoDCs producing IL-1β stimulated as in D. **G**. Percentage of Th17 cells induced upon coculture of human T cells with MoDCs from F. Coculture was maintained for 6 days under suboptimal Th17-polarizing condition in the presence of Dynabeads™ Human T-Activator CD3/CD28. Data representative of at least four independent experiments. Error bars represents ± SD. *P<0.05, **P<0.01, ***P<0.001, P<0.0001 One-way ANOVA.

### Butyrate restrains Th17 expansion and delays MM progression

*Prevotella* species produce short-chain fatty acids (SCFAs) from the fermentation of dietary fibers^40^. SCFAs act on epithelial and immune cells as histone deacetylases to reduce IL-17, TNFα and IL-6 production^41^. Interestingly, stool samples from mice treated with *P. melaninogenica* contained more acetate, buyrate and tended to be more reach in propionate than samples from *P. heparinolytica*-treated mice (Fig. 5A-C). Butyrate and butyrate-producing bacteria associate with beneficial outcomes in MM^42,43^ and potentiate ICB^44–46^. Indeed, administration of butyrate delayed disease appearance in t-Vk*MYC that was further retarded by combining anti-PD-L1 antibodies (Fig. 5D). The BM of butyrate-treated mice contained less DCs producing IL-6 (Fig. 5E) and IL-1β (Fig. 5F) and less Th17 cells (Fig. 5G) than mice treated with vehicle. Butyrate did not impact IFNγ-producing CD8^+^ T cells induced by ICB (Fig. 5H). Thus, butyrate recapitulated the *in vivo* effects of *P. melaninogenica* (Fig. 1I and Fig. S2I,J).

**Figure 5.**
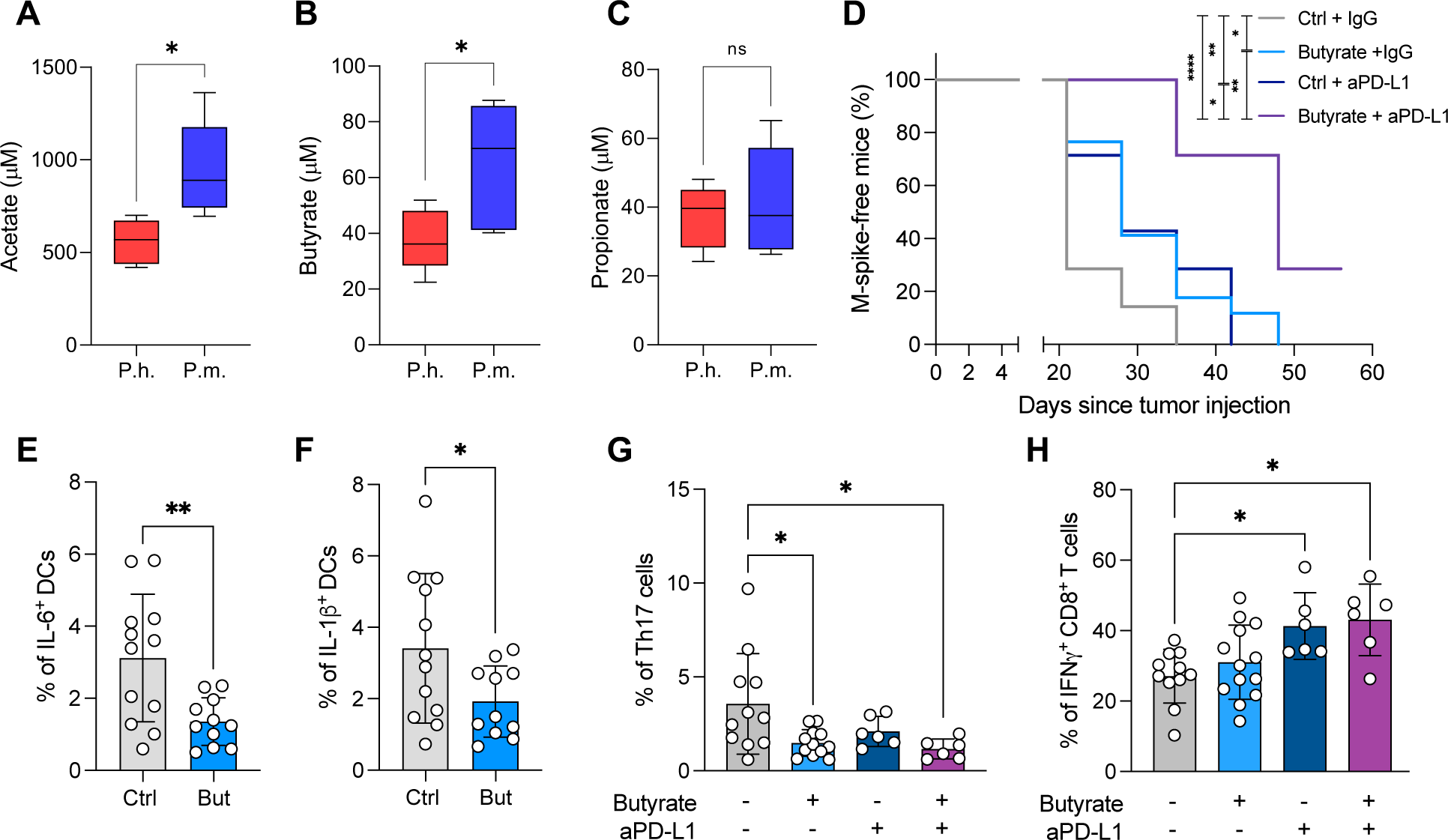
Butyrate increase by *P. melaninogenica* restrains Th17 expansion and delays MM progression. **A-B**. NMR quantification of acetate (**A)**, butyrate (**B**) and propionate (**C**) in the feces of t-Vk*MYC mice treated as in figure 4A. n=10 mice/group. Box-and-whisker plot. *P<0.05 Unpaired Student’s t-test. **D,E**. Quantification of DCs producing IL-6 (**D**) and IL-1β (E) in the BM of t-Vk*MYC mice fed butyrate (100mM in the drinking water) or vehicle (Ctrl) by flow cytometry. Butyrate treatment started two weeks before tumor inoculation. Each dot represents one mouse. Error bars represents ± SD. *P<0.05, **P<0.01. Unpaired Student’s t-test. **F**. Kaplan-Meier plot representing M-spike appearance since tumor injection of t-Vk*MYC mice fed butyrate or vehicle (Ctrl) and treated with αPD-L1 (200μg, three times/week) starting one week after tumor injection. Ctrl+IgG n=14; Butyrate+IgG n=17; Ctrl+anti-PD-L1 n=7; Butyrate+anti-PD-L1 n=7. *P<0.05. Log-rank (Mantel–Cox) test. *P<0.05, **P<0.01, ****P<0.0001. **G,H.** Percentage of Th17 cells (**G**) and IFNγ-producing CD8^+^ T cells (**H**) quantified by flow cytometry in the BM of mice treated as in **F**. Each dot represents one mouse. Data pooled from three independent experiments. Error bars represents ± SD. *P<0.05. One-Way ANOVA.

## Discussion

Clinical and experimental evidence support a role for the gut microbiome in response to immunotherapies that include ICB^5,47^. Fecal microbiota transplantation (FMT) from ICB-sensitive into ICB-refractory patients induced microbiota perturbation that correlated with enhanced anti-tumor immunity and clinical benefits^48,49^ and abrogated ICB-associated colitis^50^. Whereas several clinical trials combining FMT and ICB are underway^32^, FMT for cancer patients is far from being standardized, it does not appear to reduce incidence and severity of IrAEs^51^, may transmit multi-drug resistant bacteria, and is therefore under scrutiny by national health agencies^52^. Administration of selected bacteria is an alternative strategy to modulate the gut microbiota^12–14^. Indeed, delivery of *P. melaninogenica* to mice affected by Early-MM delayed progression to Late-MM, synergized with anti-PD-L1 antibodies and contained ICB-related skin toxicity. Thus, a strategy as simple as providing a selected commensal may intercept MM evolution.

Mechanistically, administration of *P. melaninogenica* limited the expansion of Th17 cells both in the Payer’s patches and in the BM of Vk*MYC mice. DCs were strongly affected by *Prevotellae* because *ex vivo* less DCs from the BM of *P. melaninogenica*-treated mice produced IL-6 and IL-1β when compared to mice challenged with *P. heparinolytica*. Restrained production of pro-Th17 cytokines was also obtained by challenging *in vitro* BM-derived DCs with *P. melaninogenica*. Importantly, *P. heparinolytica*-conditioned DCs generated more Th17 than *P. melaninogenica*-treated DCs. All together, these data support a model by which P*. melaninogenica* modulates the function of DCs limiting their Th17-skew potential. Because hMoDCs similarly responded *in vitro* to the two *Prevotellae*, administering *P. melaninogenica* to patients affected by plasma cell dyscrasias should be beneficial.

Our data also support the *in vivo* beneficial effects of SCFAs. Indeed, more SCFAs were found in stools of *Prevotella*-treated mice, and *P. melaninogenica* associated with much higher stool content of acetate and butyrate. Butyrate boosts the effector function of cytotoxic T lymphocytes^44^ and their memory potential^44^ buy also interferes with DC maturation^53^ and cytokine secretion^54^.Our data showing that butyrate delayed disease appearance in t-Vk*MYC mice by acting on DCs and restraining Th17 cells provide additional information on the *in vivo* beneficial effects of butyrate in in MM patients^55^.

SCFA may also be beneficial in limiting BM damage, a clinical culprit of MM^56^ because Th17 cells contributes to bone destruction^57^, and restraining Th17 expansion in MM patients by administering *P. melaninogenica* might impact bone formation. On the same line, butyrate stimulates bone formation through the secretion of Wnt10b by BM CD8^+^ T cells ^58^. Likely, the beneficial activity of *P. melaninogenica* extends beyond SCFA production.

A relevant novelty of this paper is that the combination of *P. melaninogenica* and anti-PD-L1 antibodies significantly delayed evolution to Late-MM. Importantly, *P. melaninogenica* limited the expansion of Th17 cells while fully unleashing the cytotoxic response elicited by ICB, providing treated animals with a more favorable CD8^+^ IFNγ^+^/Th17 ratio. Interestingly, IL-17 was found elevated in microsatellite stable CRC and blocking IL-17 enhanced the efficacy of anti-PD-1 therapy in mouse models of CRC^59^. These data appear at odd with those reporting that low baseline butyrate was associated with longer PFS in patients affected by metastatic melanoma^60^. IL-17 also supports the clinical benefit of dual CTLA-4 and PD-1 inhibition in melanoma patients^61^. Thus, *P. melaninogenica* and SCFA producing commensals might not be indicated when IL-17 is anti-tumoral. Conversely, *P. melaninogenica* and ICB might synergize in patients affected by other hematologic and solid tumors in which IL-17 acts as driving force^11^.

We also found that modulation of the gut microbioma with SCFA-producing bacteria limits IrAEs. This effects again appears related to the ability of *P. melaninogenica* to limit Th17 cell expansion especially in skin-draining lymph nodes. Interestingly, mIHC analyses of the skin from mice receiving imiquimod, anti-PD-L1 antibodies and *P. melaninogenica* showed reduced density of gut-homing (i.e., α4β7^+^) cells, supporting restrained migration of cells from the gut into the skin of mice receiving *P. melaninogenica*. CD11c^+^ and CD11c^+^IFNγ^+^ cells were also less represented and tended to be more distant from CD4 and CD8 T cells in *P. melaninogenica*-treated mice, suggesting that the intestinal commensal in this model of skin toxicity also impacted APCs.

IL-17 is involved in several IrAEs^62^, ICB may unleash commensal-specific Th17 responses leading to skin toxicity^23^, and IL-17 blockade reversed psoriatic rash in one patient treated with pembrolizumab^63^. Inhibition of IL-17 also abrogated nivolumab-associated colitis, and protected mice against ICB-induced thyroiditis ^26^. ICB-induced colitis can be ameliorated by treating mice with *Bifidobacterium*, which favors the expansion of Tregs^64^, a mechanism distinct form *P. melaninogenica*. Thus, *P. melaninogenica*, by producing SCFAs and restraining Th17 cell expansion might positively impact additional IrAEs.

As discussed above, microbiota perturbation holds safety issues. Abundance of *P. melaninogenica* together with other commensals were found in the lung of patients affected by cystic fibrosis^65^, in the oral cavity of children with halitosis^66^ and active caries^67^ and in the gut of patients affected by ankylosing spondylitis^68^. These data suggest that in selected clinical conditions, *P. melaninogenica* may act as pathobiont. While daily administration of *P. melaninogenica* to mice promoted gastric inflammation^69^, intranasal administration of *P. melaninogenica* did not cause any damage to healthy mice, but it enhanced bleomycin-induced lung immune infiltration and fibrosis^70^. Conversely, *P. melaninogenica* suppressed experimental autoimmune encephalomyelitis in HLA-DR3.DQ8 transgenic mice^71^. Different outcomes may relate to different experimental settings and/or strains of *P. melaninogenica* as reported for *P. copri*^72^. We also cannot exclude that *P. melaninogenica* drove dissimilar reconstitution of the microbiota community in the various experimental settings described above.

Altogether, our results support *P. melaninogenica* administration as a strategy sufficient *per se* to modify the trajectory of patients affected by SMM. The combination of *P. melaninogenica* and ICB might also represent an additional strategy to uncouple efficacy and toxicity of ICB and limit MM aggressiveness.

## Acknowledgments

This work was supported Leukemia and Lymphoma Society (L&LS; Grant #6618-21 to MB) and the Cancer Research Institute (CLIP Grant #CRI4838 to M.B.). The research leading to these results has also received funding from AIRC under IG 2018 -ID. 21808 project – P.I. M.B. L.L.C. conducted this study in partial fulfillment of her Ph.D. at San Raffaele University. We thank the International Myeloma Society (IMS) for the IMS Young Investigation Award to L.L.C. We thank Dr. Arianna Brevi for the initial experimental training. We thank Dr. Valeria Mannella from Proteomics and Metabolomics Facility (ProMeFa; IRCCS Ospedale San Raffaele). We thank Dr. Amleto Fiocchi from the Animal Histopathology facility (IRCCS Ospedale San Raffaele and Universita Vita-Salute San Raffaele).

## Authorship Contributions

M.B. and L.L.C. designed and conceived the study. L.L.C. performed most of the experiments and analyzed data. P.Z. and B.M. performed *in vitro* experiments with human cells. M.G. helped with *in vivo* experiments and maintained the murine colonies. A.P., M.L. and G.M. helped with some *in vivo* experiments. N.R. and F.S. performed histological evaluation and scoring. S.S. and N.C. provided and maintained bacteria cultures. A.T. and A.R. performed and analyzed Multiplex IHC. M.C. and L.B. provided genetically engineered mice. L.L.C. prepared the figures and tables. M.B and L.L.C. wrote the manuscript. All authors contributed to critically reading the manuscript and approved the final version.

