## Supplemental Data for "Microbiome Modulation Uncouples Efficacy and Toxicity Induced by Programmed Death-1/Programmed Death-Ligand1 Blockade"

^1^Cellular immunology Unit, Division of Immunology, Transplantation and Infectious Diseases, IRCCS Ospedale San Raffaele, Milan, Italy; ^2^Vita-Salute San Raffaele University, Milan, Italy; ^3^Immunology and Molecular Oncology Diagnostics, Veneto Institute of Oncology IOV-IRCCS, Padua, Italy; ^4^Pathology and histology department, IRCCS Ospedale San Raffaele, Milan, Italy; ^5^Department of Surgery, Oncology and Gastroenterology (DiSCOG), University of Padua, Padua, Italy; ^6^Laboratory of Microbiology, IRCCS Ospedale San Raffaele, Milan, Italy; ^7^Mayo Clinic, Arizona, USA.

**Supplemental information**

Supplemental Methods

Supplemental Table 1

Supplemental Table 2

Supplemental Figure 1

Supplemental Figure 2

Supplemental Figure 3

**Supplemental methods:**

**Serum protein electrophoresis.** Mouse blood was periodically collected. Semi-automated electrophoresis was performed on the Hydrasys instrument (Sebia, Lissex, France). According to the manufacturer’s instructions, 10 μL of undiluted serum were manually applied to the Hydragel agarose gels (Sebia). The subsequent steps: electrophoresis (pH 9.2, 20W constant current at 20 °C), drying, amidoblack staining, de-staining and final drying were carried out automatically. The use of Hydrasys densitometer and Phoresis software (Sebia) for scanning resulting profiles provided accurate relative concentrations (percentage) of individual protein zones. M-spike levels were calculated as total gamma globulins/albumin ratio (γ/albumin).

**Isolation of single cell suspension and flow cytometry**. Mouse bones were harvested, and epiphyses were cut to collect BM cells. Peyer’s Patches were removed from the Small Intestine of the same animals and gently disaggregated with the help of tweezers. Single cell suspensions were labeled with fluorochrome-conjugated monoclonal antibodies (Table S1) and acquired by BD FACSCanto™ II, BD FACSymphony™ A5 (BD Bioscience) or CytoFLEX LX (Beckman Coulter). Cells were also assessed for intracellular cytokine production after stimulation with Phorbol Myristate Acetate (PMA) (1mg/mL; MedChemExpress) and ionomycin (1mg/mL; MedChemExpress) in the presence of Brefeldin A (BFA; 10µg/mL, MedChemExpress). After incubation, cells were washed and stained for surface markers, fixed and permeabilized with Fixation/Permeabilization Kit (BD-Bioscience) or with paraformaldehyde 2% and saponin. Cells were then washed and stained for intracellular markers. Data were analyzed using the FlowJo software (Treestar Inc).

**Multiplex immunohistochemistry imaging and analysis**. Multiplex stained slides were imaged using the Mantra Quantitative Pathology Workstation (Akoya Biosciences, Marlborough, MA, USA) at 20X magnification. The inForm Image Analysis software (version 2.4.9, Akoya Biosciences, Marlborough, MA, USA) was used to unmix multispectral images. A selection of representative multispectral images was used to train the inForm software to create algorithms to apply in the batch analysis of all acquired multispectral images. phenoptrReports (add-ins for R Studio from Akoya Biosciences) was used to calculate cell density and spatial metrics. Cell density data were calculated as the sum of the cells positive for a specific marker, divided by the area analyzed from the same tissue slide. For mean distance between different cell subtypes, the nearest neighbour analysis was used.

**Supplemental Tables**

**Table S1**

| Reactivity | Marker | Fluorochrome | Clone | Provider |
| --- | --- | --- | --- | --- |
| anti-mouse | CD45 | V500 | 30-F11 | BD Biosciences |
| anti-mouse | CD45 | BUV395 | 30-F11 | BD Biosciences |
| anti-mouse | CD3 | FITC | 145-2C11 | Biolegend |
| anti-mouse | CD3 | BUV737 | 145-2C11 | BD Biosciences |
| anti-mouse | CD3 | APC-Cy7 | 145-2C11 | Biolegend |
| anti-mouse | CD4 | PerCP-Cy5.5 | RM4-5 | Biolegend |
| anti-mouse | CD4 | BV605 | RM4-5 | Biolegend |
| anti-mouse | CD8 | FITC | 53-6.7 | Biolegend |
| anti-mouse | CD8 | V500 | 53-6.7 | BD Biosciences |
| anti-mouse | CD8 | PE-Cy7 | 53-6.7 | Biolegend |
| anti-mouse | CD8 | BV650 | 53-6.7 | Biolegend |
| anti-mouse | CD25 | APC | PC61 | Biolegend |
| anti-mouse | B220 | FITC | RA3-6B2 | Biolegend |
| anti-mouse | CD138 | PE | 281-2 | BD Biosciences |
| anti-mouse | IA^b^ | FITC | 25-9-17 | Biolegend |
| anti-mouse | IA^b^ | PE | AF6-120.1 | BD Biosciences |
| anti-mouse | CD11b | Pacific-blue | M1/70 | Biolegend |
| anti-mouse | CD11c | PE-Cy7 | N418 | Biolegend |
| anti-mouse | PD-L1 | BV421 | MIH5 | BD Biosciences |
| anti-mouse | IL-17 | PE | TC11-18H10 | BD Biosciences |
| anti-mouse | IFN𝛾 | Alexa Fluor 700 | XMG1.2 | Biolegend |
| anti-mouse | IFN𝛾 | APC | XMG1.2 | Biolegend |
| anti-mouse | IL-6 | APC | MP5-20F3 | Biolegend |
| anti-mouse | IL-1β | APC Vio770 | REA577 | Miltenyi Biotec |
| anti-mouse | FOXP3 | PE | FJK-16s | eBioscience |
| anti-human | CD11c | Pe-Dazzle | 3.9 | Biolegend |
| anti-human | CD3 | FITC | HIT3a | Biolegend |
| anti-human | CD4 | APC-Cy7 | RPA-T4 | BD Biosciences |
| anti-human | CD8 | BUV-805 | SK1 | BD Biosciences |
| anti-human | IL-17 | BV-711 | BL168 | Biolegend |
| anti-human | IFN𝛾 | BUV-395 | B27 | BD Biosciences |
| anti-human | IL-1β | FITC | [JK1B-1](https://www.biolegend.com/en-ie/search-results?Clone=JK1B-1) | eBioscience |
| Live/dead markers | | |  |  |
| Zombie-Aqua^TM^ | | |  | Biolegend |
| Viakrome-808 | | |  | Beckman Coulter |

**Table S1.** Mouse and human fluorochrome-conjugated monoclonal antibodies for flow cytometry.

**Table S2**

| Host | Reactivity | Marker | Clone | Provider |
| --- | --- | --- | --- | --- |
| rabbit | anti-mouse | IFN-γ | polyclonal | Novus Biologicals |
| rat | anti-mouse | IL-17A | TC11-18H10 | Novus Biologicals |
| rat | anti-mouse | integrin ⍺4β7 | RM0059-2G18 | Abcam |
| rabbit | anti-mouse | CD11c | D1V9Y | Cell Signalling |
| rat | anti-mouse | CD4 | 4SM95 | Thermo Fisher |
| rat | anti-mouse | CD8a | 4SM15 | Thermo Fisher |
| rat | anti-mouse | FoxP3 | FJK-16S | Thermo Fisher |
| rabbit | anti-mouse | Cytokeratin 19 | 10712-1-AP | Proteintech |

**Table S2.** Multiplex immunohistochemistry antibody panel.

**Supplemental Figures**

**Figure S1**

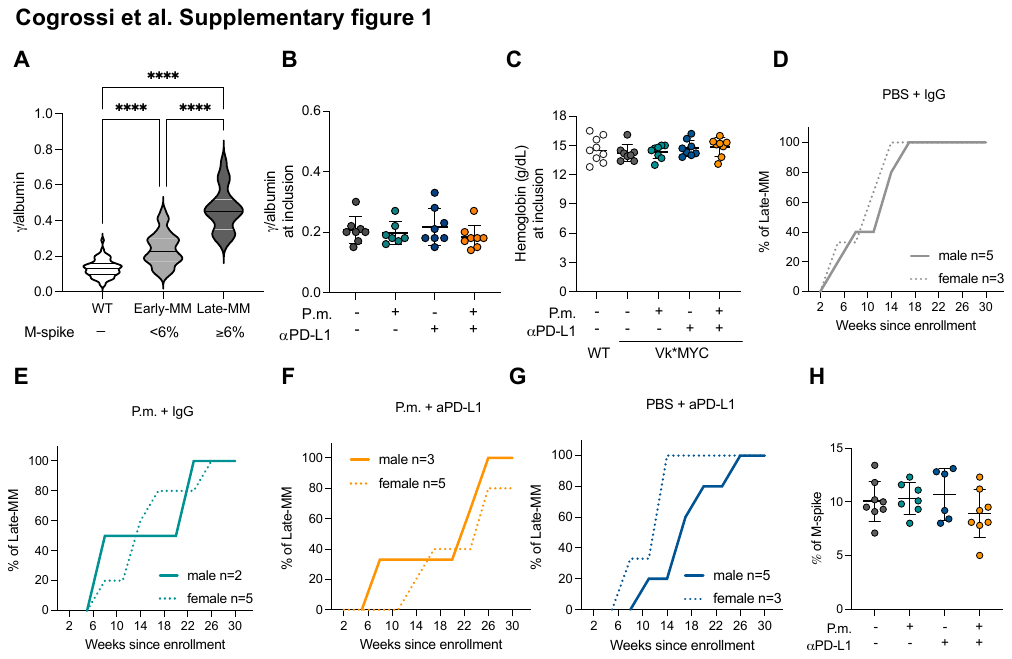

**Figure S1. A.** Quantification of M-spike reported as γ/albumin ratio for wild-type mice (WT; n=62), Early-MM (n=65), and Late-MM (n=22) Vk*MYC mice. ****P<0.0001 One-Way ANOVA. **B.** γ/albumin ratio for each Early-MM Vk*MYC mouse at the time of enrollment. Error bars represent ± SD. **C**. Quantification of hemoglobin from the peripheral blood of each Early-Vk*MYC mouse at enrollment. WT: wild-type age-matched controls. Error bars represent ± SD. **D-G**. Percentage of Vk*MYC mice treated with PBS+IgG (**B**), P.m.+IgG (**C**), PBS+aPD-L1 **(D**), P.m.+aPD-L1 (**E**) progressing to Late-MM segregated for sex. **H**. M-spike at sacrifice of Vk*MYC mice from Fig. 1A-E.

**Figure S2**

IL-17

Bone marrow

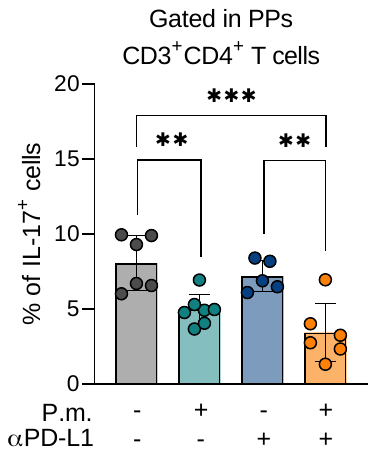

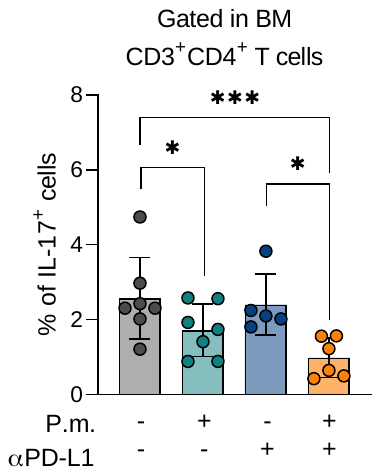

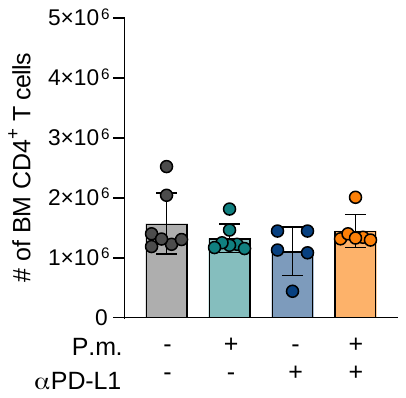

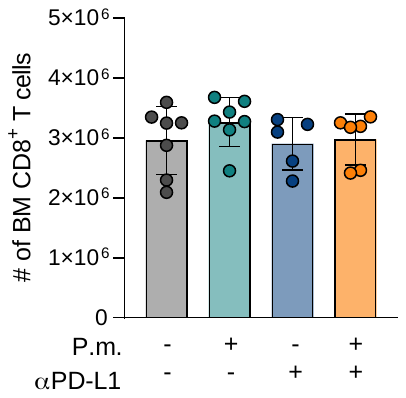

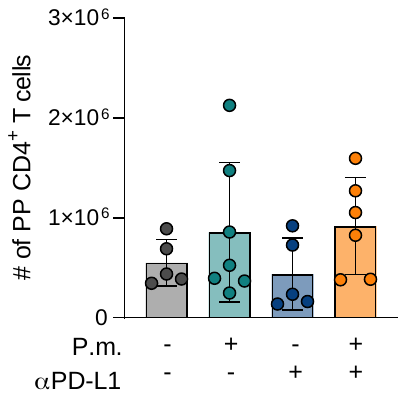

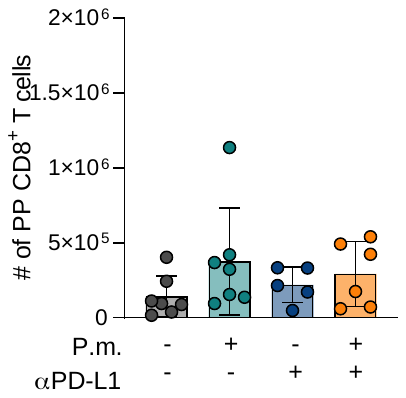

**A**

**B**

**C**

**D**

**E**

**F**

**G**

**H**

**I**

**K**

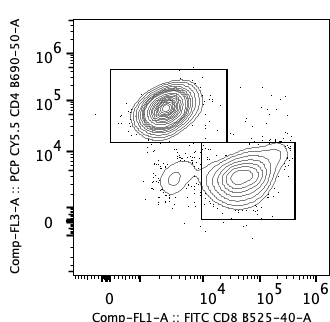

CD4

CD8

Peyer’s Patches

CD4

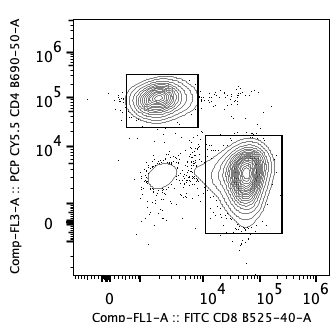

CD8

Peyer’s Patches

Bone marrow

**J**

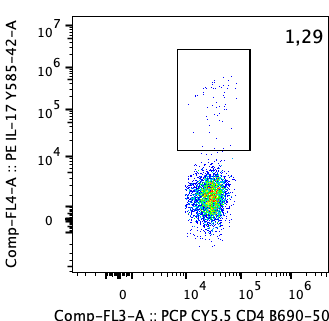

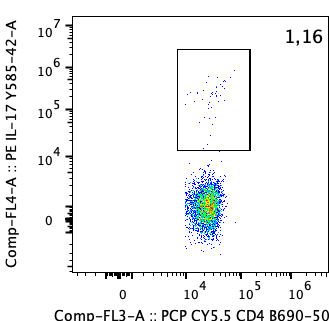

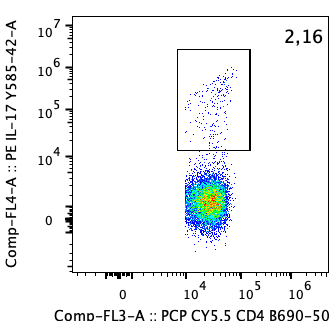

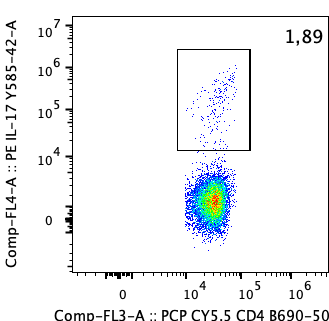

αPD-L1

IgG

PBS

P.m.

CD4

**N**

**L**

**M**

IgG

IL-17

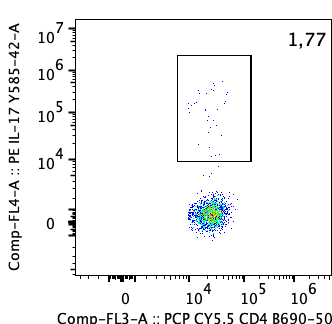

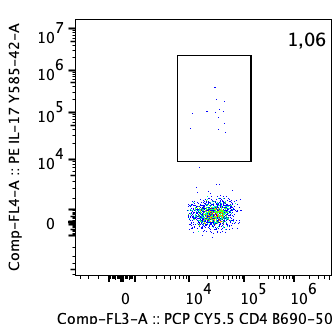

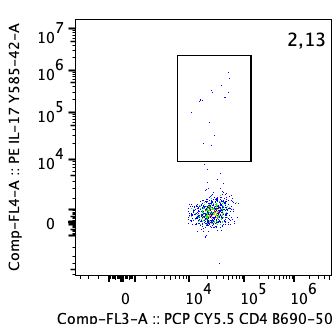

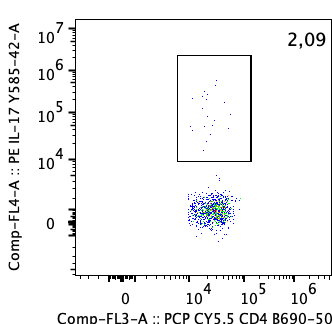

αPD-L1

PBS

P.m.

CD4

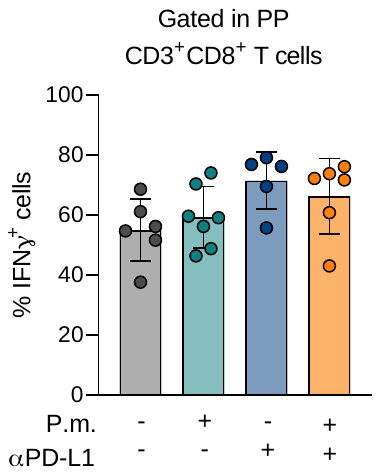

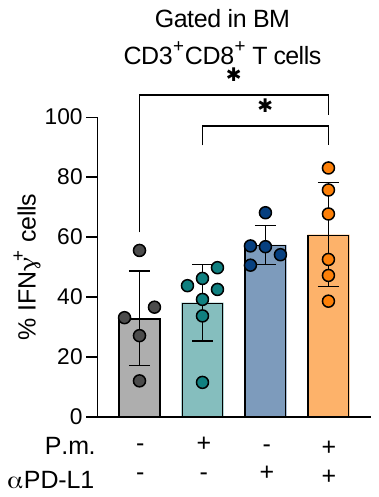

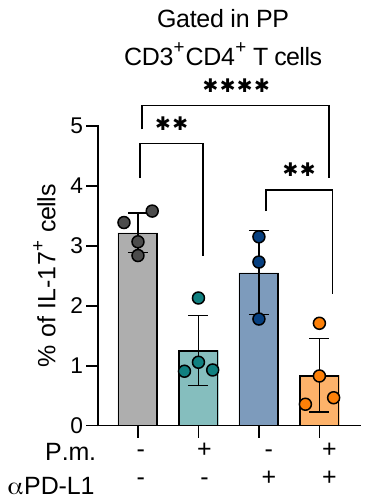

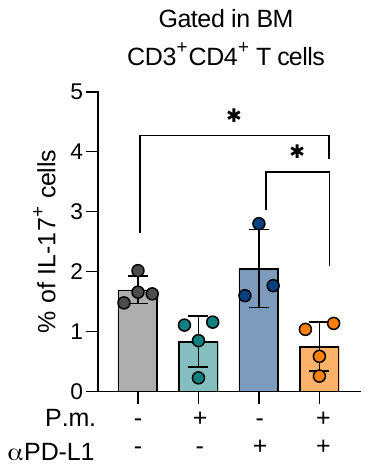

**Figure S2. A-J. Flow cytometry data from mice reported in figure 2**. Representative plot (**A**) and quantification of CD4^+^ (**B**) and CD8^+^ (**C**) T cells in the intestinal PPs. **D**. Percentage of IL-17^+^ cells gated in CD4^+^ T cells in the PPs. **E**. Percentage of IFNγ^+^ cells gated in CD8^+^ T cells in the PPs. Representative plot **(F)** and quantification of CD4^+^ (**G**) and CD8^+^ (**H**) T cells in the BM. **I**. Percentage of IL-17^+^ cells gated in CD4^+^ T cells in the BM. **J.** Percentage of IFNγ^+^ cells gated in CD8^+^ T cells in the BM. **K-N**. Flow cytometry data from t-Vk*MYC mice described in figure 1G-I. Representative plot (**K**) and percentage (**L**) of Th17 in the PPs. Representative plot (**M**) and percentage (**N**) of Th17 in the BM. Data are gated in CD4^+^ T cells. Each dot is one mouse. Error bars represent ± SD. *P<0.05, **P<0.01, ***P<0.001, ****P<0.0001 One-Way ANOVA.

**Figure S3**

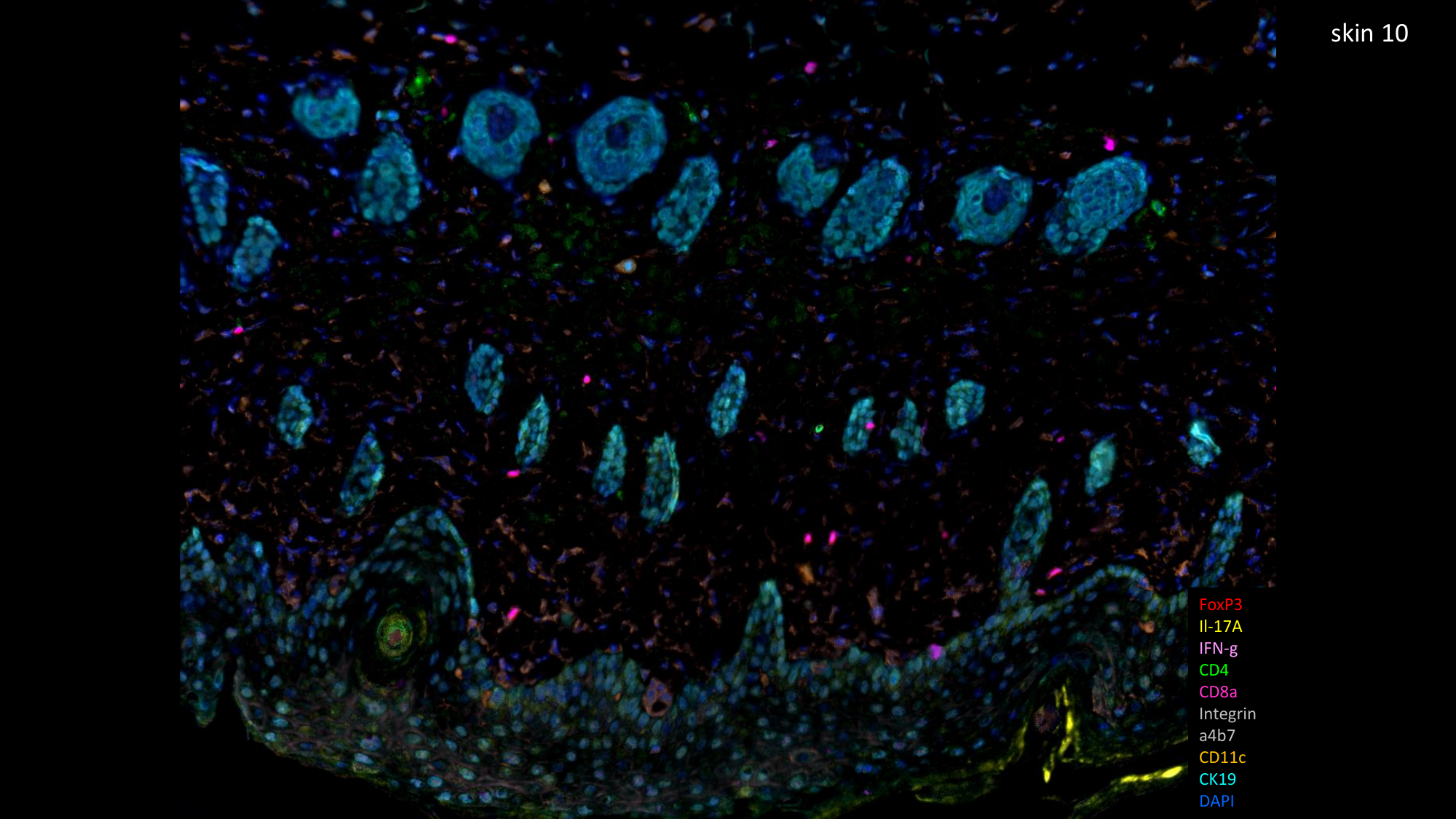

IMQ+ αPD-L1+P.m*.*

IMQ+αPD-L1

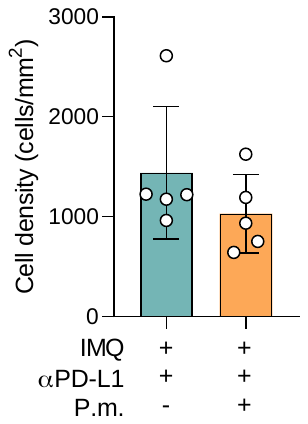

IL-17^+^

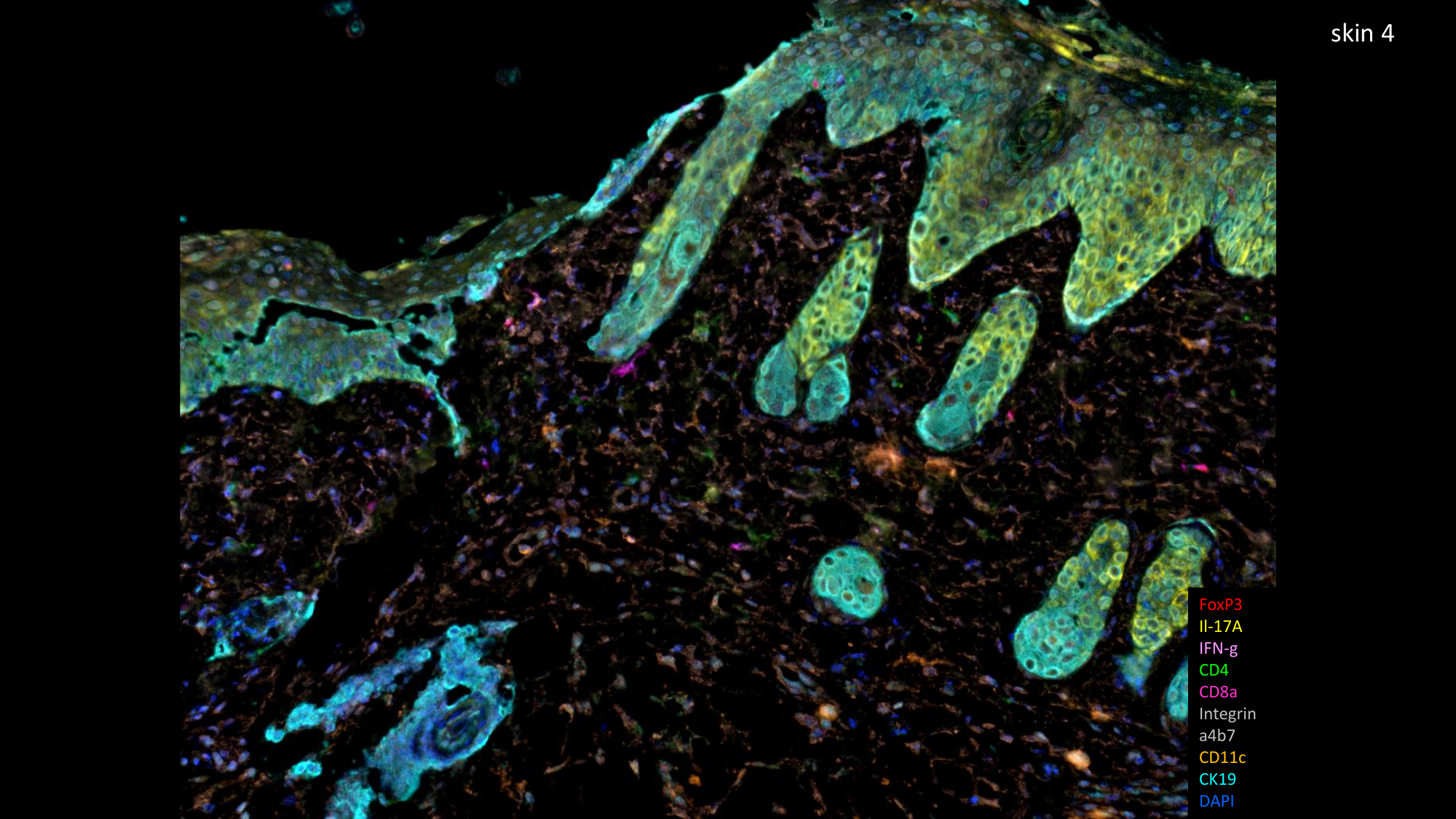

**A**

**B**

**C**

**D**

CD11c^+^ to CD4^+^

CD11c^+^ to CD8^+^

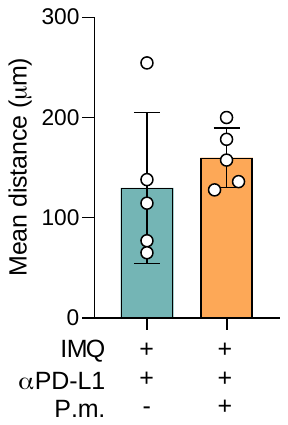

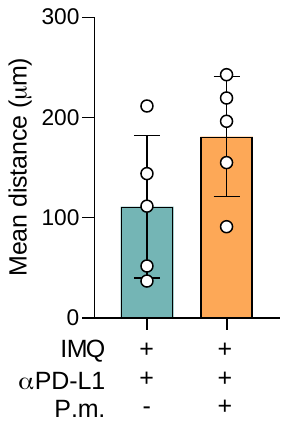

**Figure S3. A.** Representative multiplex IHC images from mice treated with IMQ + αPD-L1 in the absence (left) or presence (right) of P.m*.* administration. Magnification 20X. **B**. Cell density per slide of IL-17^+^ cells. **C-D**. Mean distance between CD11c^+^ and CD4^+^ T cells (**C**) or CD8^+^ T cells (**D**) per slide.
